# DNA supercoiling enhances DNA condensation by ParB proteins

**DOI:** 10.1101/2024.07.11.603076

**Authors:** Alejandro Martin Gonzalez, Miloš Tišma, Brian T. Analikwu, Anders Barth, Richard Janissen, Hammam Antar, Gianluca Kemps, Stephan Gruber, Cees Dekker

**Author notes:** These authors have contributed equally (Alejandro Martin Gonzalez, Miloš Tišma).

## Abstract

The ParABS system plays a critical role in bacterial chromosome segregation. The key component of this system, ParB, loads and spreads along DNA to form a local protein-DNA condensate known as a partition complex. As bacterial chromosomes are heavily supercoiled due to the continuous action of RNA polymerases, gyrases, and nucleoid-associated proteins, it is important to study the impact of DNA supercoiling on the ParB-DNA partition complex formation. Here, we use an *in vitro* single-molecule assay to visualize ParB on supercoiled DNA. Unlike most DNA-binding proteins, individual ParB proteins are found to not pin plectonemes on supercoiled DNA, but freely diffuse along supercoiled DNA. We find that DNA supercoiling enhances ParB-DNA condensation which initiates at lower ParB concentrations than on DNA that is torsionally relaxed. ParB proteins induce a DNA-protein condensate that strikingly absorbs all supercoiling writhe. Our findings provide mechanistic insights that have important implications for our understanding of bacterial chromosome organization and segregation.

## Introduction

Reliable segregation of chromosomes to daughter cells is a fundamental requirement for the stable propagation of all living organisms. The ParABS system is the primary mechanism responsible for the faithful segregation of chromosomes in the majority of bacteria^1,2^. It is comprised of an ATP-hydrolase partition protein A (ParA)^3,4^, a CTP-hydrolase partition protein B (ParB) ^5,6^, and a 16-base pair centromeric sequence known as *parS*^7^ that is present in multiple copies near the origin of replication^7^. While ParA proteins bind non-specifically to the DNA to cover the entire chromosome^8,9^, ParB proteins specifically load onto DNA at the *parS* sequence^6,10,11^, whereupon they diffuse away along the DNA to subsequently use dynamic ParB-ParB bridging ^12–15^ to assemble into a higher-order nucleoprotein structure known as a “partition complex” ^16–18^. The partition complex promotes the loading of SMC proteins^19–21^ and interacts with a ParA gradient along the nucleoid^22–25^ which leads to the accurate segregation of the nascent origins and chromosomes to daughter cells.

In cells, transcription exerts prominent forces and twists on the genomic DNA^26,27^. An RNA polymerase transcribing a gene will continuously create positive supercoiling downstream and negative supercoiling upstream of the transcription site^28–31^. When such torsional tension is built up, the DNA will twist and form plectonemes which are extended intertwined DNA helices^31–33^. Many other cellular processes modulate and regulate the supercoiling within the genome through actions of topoisomerases, gyrases, and nucleoid-associated proteins (NAPs). Vice versa, supercoiling is known to affect many processes in bacterial cells by changing the protein binding affinities and spatial organization of the DNA^26,34^. Recent *in silico* studies^35,36^ suggested that supercoiling may, for example, strongly change the dynamics of the ParB-DNA partition complex by modulating the interactions between distal segments of the DNA.

Here, we examine the effects of DNA supercoiling on ParB diffusion and condensation dynamics using a single-molecule visualization assay^37^. We measured the diffusion of single ParB dimers on supercoiled and non-coiled DNA molecules, and found that the presence of supercoiling reduces the residence time of ParB on the DNA. Furthermore, we measured the degree of ParB-DNA condensation induced by ParB proteins on supercoiled DNA as well as the changes in plectoneme dynamics and localization in the presence of ParB. We observed that all plectonemic structures were absorbed within the ParB-DNA condensate. Lateral DNA flow and atomic force microscopy (AFM) confirmed that supercoiled DNA in the presence of ParB yielded a collapsed structure that absorbed all supercoiling writhe. This experimental study reveals the interplay of DNA supercoiling and ParB-DNA condensation, two fundamental and essential processes that govern bacterial genome organization and segregation.

## Results

To observe ParB binding to supercoiled DNA molecules, we employed a single-molecule DNA stretching assay^38^ with fluorescently labelled ParB^alexa647^ proteins^39^. We attached a 38 kb DNA molecule that contained a *parS* site close to the middle (DNA*_parS_*) to a glass surface via multiple biotin-streptavidin interactions at both DNA ends (Fig. 1A, see Methods for details). Due to the multiple biotin attachment points at each end, a large fraction of such DNA*_parS_*molecules was torsionally constrained and could not rotate around its central axis to relieve any torsional strain on the molecule. We exploited this feature to directly introduce supercoiling of the desired handedness, either positive or negative supercoiling, to the DNA molecule. We achieved this by changing the concentrations of the intercalator dye SYTOX Orange (SxO) that is also used for fluorescent visualization of the DNA^40–42^ (Fig. 1B, see Methods for details). When a SxO fluorophore intercalates into the dsDNA helix, it locally pushes two base-pairs apart and thus, due to the induced change of the base-pair distance and angle^43,44^, locally underwinds the DNA which yields an overwinding twist into the remainder of the molecule since the linking number of the molecule is fixed due to the tethering. For sufficiently large SxO concentration, this overwinding yields a positive writhe in the DNA molecule, i.e. positively coiled plectonemes (Fig.1B). The latter were observed as dynamically moving local high-density spots on the linearly stretched DNA molecule^38^ (Fig. 1E; Movie S1). The effect of supercoiling on ParB binding and condensation was monitored for both supercoiled and torsionally unconstrained DNA molecules in the same field of view^42^ (Fig. S1).

**Figure 1.**
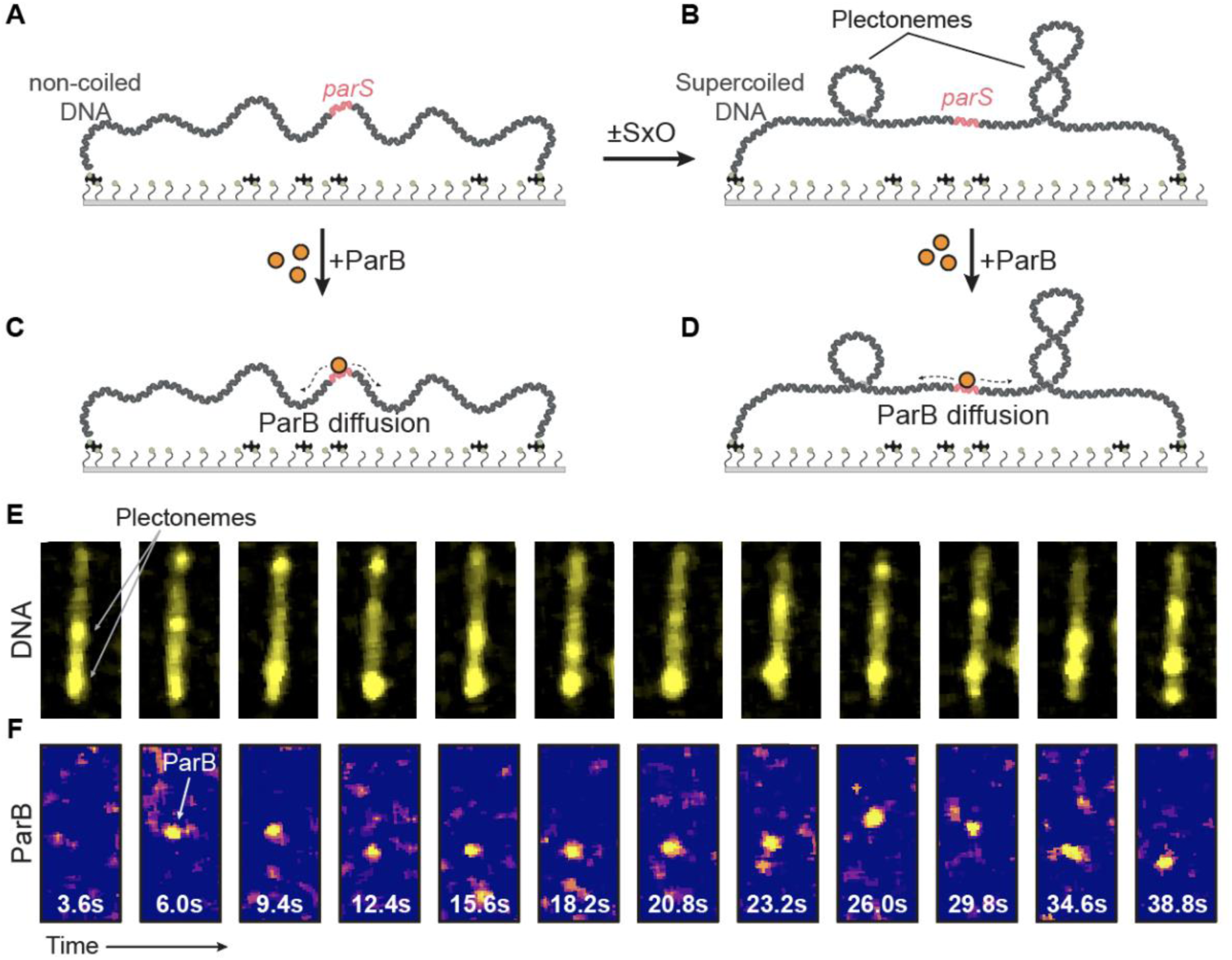
*In vitro* single-molecule fluorescence for studying ParB proteins on supercoiled DNA. A) Schematic representation of the single-molecule DNA stretching assay, with 38 kb DNA*_parS_* tethered to the glass surface. B) The same molecule after the addition or reduction of SYTOX Orange intercalating dye which induces supercoiling and plectoneme formation. C-D) same as AB but with ParB added to the nicked or supercoiled DNA molecules, respectively. E) Images of DNA molecules taken at various times, showing dynamic plectoneme movement on the supercoiled DNA molecule. F) Visualization of single ParB^alexa647^ dimer on the DNA molecule of panel E, showing binding and one- dimensional diffusion along the DNA*_parS_*.

### ParB proteins efficiently bind and diffuse on supercoiled DNA

After forming positively supercoiled DNA, we added ParB^alexa647^ (Fig. 1C, D) to observe protein localization and their diffusion behavior along the DNA, similar to our previous report for torsionally unconstrained DNA^39^. ParB proteins were observed to load onto the DNA at the *parS* site and to immediately exhibit one-dimensional diffusion away from the binding site, as best observed in a kymograph (Fig. S2). While ParB diffusion was shown previously on non-coiled DNA molecules^12,39,45^, we here observed the efficient loading and diffusion of ParB proteins along the supercoiled DNA molecules. Unexpectedly, we observed that ParB did *not* appear to strongly pin plectonemes (Fig. 1E, F). After binding the ParB, the dynamics of the plectonemes continued in an unperturbed way, and no clear co-localization of ParB and the plectonemes was observed (Fig. 2A, B, S2B-F). This contrasts many other DNA-binding proteins that were shown to induce a localization of plectonemes at the binding site of the protein^31,46–52^, presumably because the binding induced a local change in DNA curvature that lowers the energy of plectoneme formation. We, however, observed continuous one-dimensional diffusion by ParB. ParB dimers that were diffusing on positively supercoiled DNA showed a somewhat higher diffusion coefficient D = 0.69 ± 0.35 kb^2^/s (median ± SE; n = 66, Fig. S3A-B, Table 1) than on non-coiled DNA molecules (D = 0.43 ± 0.13 kb^2^/s, median ± SE, n = 58, Fig. S3C-D).

**Figure 2.**
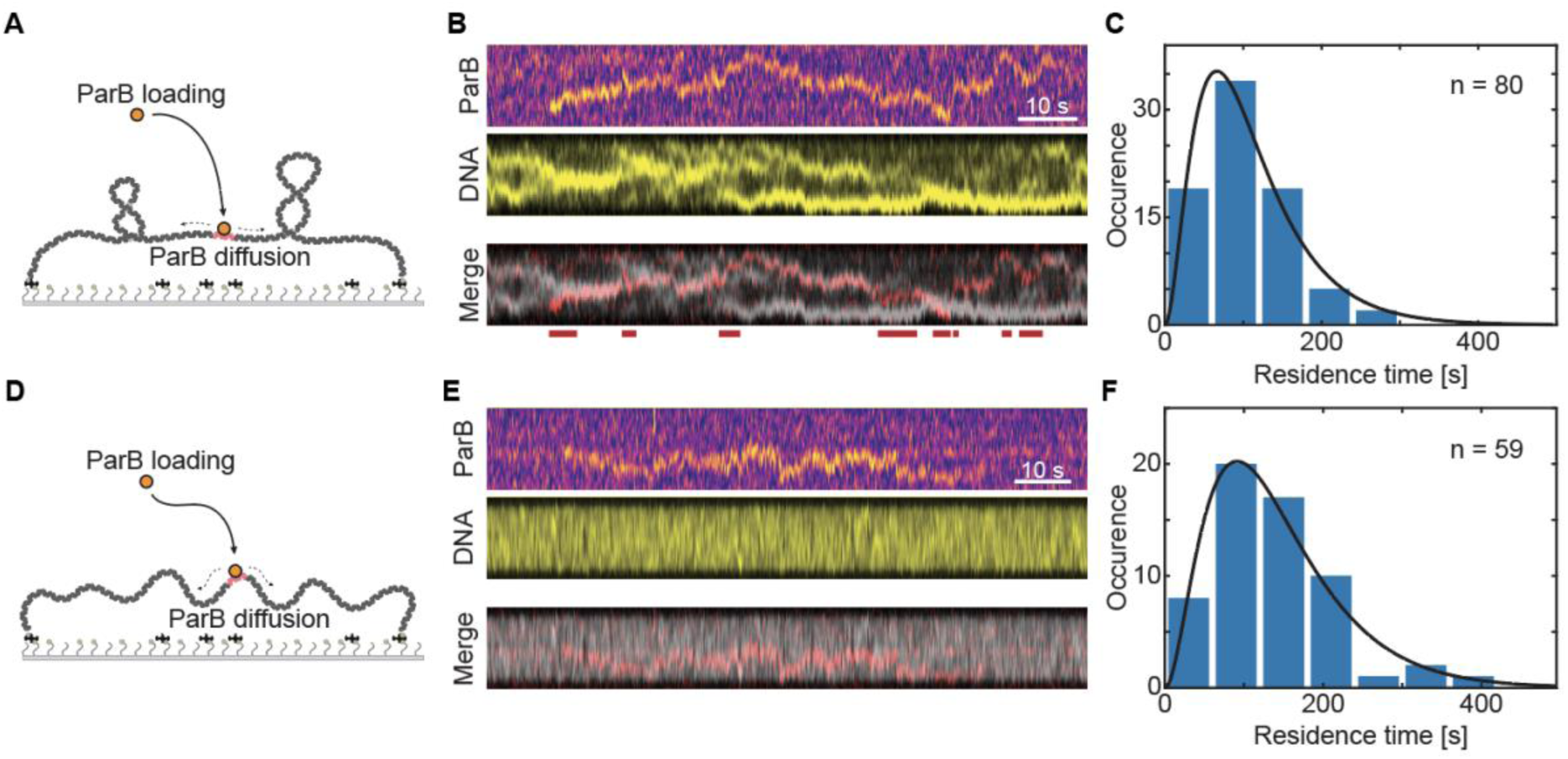
ParB efficiently binds and diffuses along the supercoiled DNA. **A)** Schematic representation of the supercoiled DNA and addition of ParB proteins. **B)** Kymographs showing one- dimensional diffusion of a single ParB^alexa647^ dimers (top) and DNA plectonemes (middle). Red lines below the merged kymograph indicate sections where ParB signal does not overlap the plectonemes. **C)** Residence times of diffusing ParB dimers after binding to the *parS* site. The data were fitted to a model assuming a delayed dissociation of ParB from the DNA after CTP hydrolysis of both nucleotides (black line, see Tišma et al. 2022). **D-F)** Same as A-C for non-coiled DNA molecules.

**Table 1.** Quantification of ParB residence times in the presence of DNA supercoiling. Significance tests compared to the non-coiled samples: MW – *Mann-Whitney test*, KS - *Kolmogorov-Smirnov test*.

| | Mean | SEM | Median | Mode | $k_{CTP}$ | $k_{off}$ | N | Significance | |
| --- | --- | --- | --- | --- | --- | --- | --- | --- | --- |
| Non-coiled | 141 s | 10 s | 129 s | 91 s | $0.016 \text{ s}^{-1}$ | $0.020 \pm 0.009 \text{ s}^{-1}$ | 59 | MW | KS |
| Positive supercoiled | 109 s | 8 s | 95 s | 70 s | $0.017 \text{ s}^{-1}$ | $0.047 \pm 0.014 \text{ s}^{-1}$ | 80 | 0.0028 | 0.0052 |
| Negative supercoiled | 106 s | 8 s | 100 s | 66 s | $0.021 \text{ s}^{-1}$ | $0.028 \pm 0.016 \text{ s}^{-1}$ | 58 | 0.0026 | 0.0072 |

We observed a non-exponential distribution for the residence time of ParB molecules on DNA (Fig. 2C), which contrasts the typical exponential decay for most DNA-binding proteins with the dissociation rate^53,54^. This indicates the existence of multiple rate-limiting steps. For ParB, both CTP molecules that are sandwiched between the ParB monomers likely need to be hydrolyzed in order to open and detach the dimer from the DNA^39,55^. To describe prolonged diffusion on the DNA molecule, we applied our previously described model^39^ which incorporates CTP hydrolysis as the rate-limiting step that extends the ParB diffusion and spreading on the DNA. From the model we obtained the residence time of 70 s (mode of distribution, Fig. 2C, Table 1) of ParB molecules diffusing on supercoiled DNA which was slightly lower in comparison to the molecules diffusing on non-coiled DNA (91 s, p-value <0.005, Fig. 2F, Table 1).

While all the data presented above were for positively supercoiled DNA, we observed very similar behavior of ParB proteins on negatively supercoiled DNA. We introduced negative supercoiling into the DNA*_parS_* molecules in the same manner described previously, except that we lowered rather than increased the dye concentration after DNA-binding (see Methods). Subsequent addition of ParB^alexa647^ molecules showed efficient binding of ParB to negatively supercoiled DNA molecules (Fig. S4A-B), with a typical residence time of 66 s (mode of distribution, Fig. S4C, Table 1), and a diffusion coefficient of 0.59 ± 0.25 kb^2^/s (median ± SE, Fig. S4E). The results were thus qualitatively and quantitatively similar for positive and negative supercoiling. Overall, the presence of DNA supercoiling did not hinder the loading of single ParB molecules, while it decreased the residence time and increased the apparent diffusion coefficient.

### Supercoiling facilitates DNA condensation by ParB proteins

In previous *in vivo* and *in vitro* studies, ParB proteins were shown to be proficient at forming DNA condensates around the *parS* site^15,16,18,45,56,57^, and various current models suggest a strong dependence on ParB diffusion and self-self interaction. Notably, distant-site binding due to supercoiling-related proximity could severely alter the formation of ParB-DNA condensates, such as was proposed recently by Connolley *et al*^35^.

We therefore set out to test the DNA condensation in the presence of DNA supercoiling at higher concentrations of ParB proteins. For comparison, we added either low (3 nM, Fig. 3A-D) or high ParB concentrations (25 nM, Fig. 3E-H) onto supercoiled DNA*_parS_* molecules. At low ParB concentration, the rapidly moving plectonemes on the DNA largely remained unaffected after ParB addition, both in their position (Fig. 3A) and the number of co-existing plectonemes formed on the supercoiled DNA (Fig. 3B, Fig. S4A-F). There also was no significant change in the amount of DNA within each plectoneme, and the total amount of DNA integrated over plectonemes remained at the same value, ∼12 kb for this set amount of supercoiling (Fig. 3C, D). Furthermore, the dynamic behavior of DNA in the presence of low ParB concentration was of comparable dynamics and localization to the control in the absence of the protein (Fig. S4G, H). Small ParB clusters were found to not correlate with the position of the plectonemes on the DNA over time, further confirming the absence of significant plectoneme pinning (Fig. S4F).

**Figure 3.**
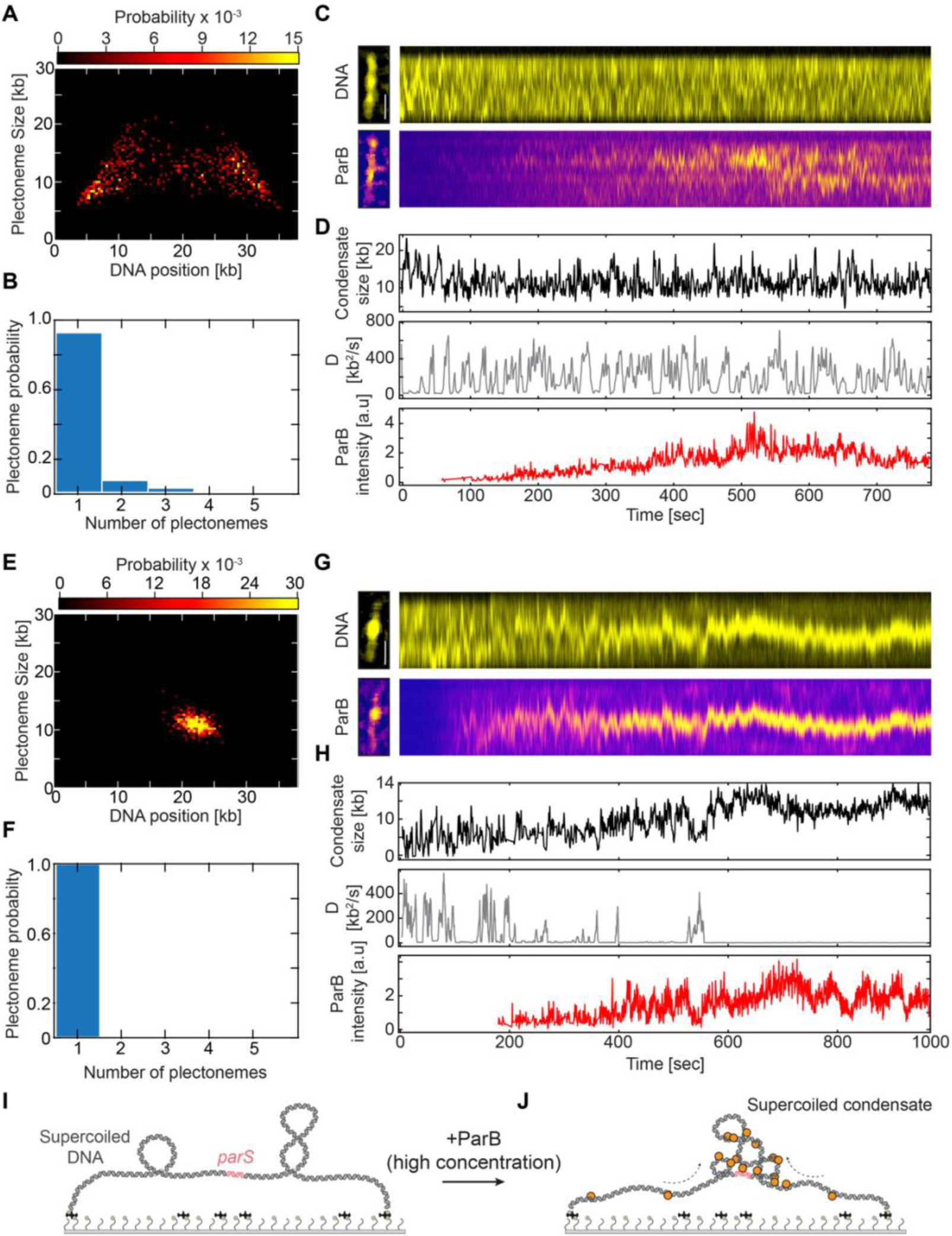
Multiple ParB proteins pin DNA plectonemes into a single static cluster. **A)** Probability distribution of plectoneme size versus DNA position at 3 nM ParB protein. **B)** Observed number of plectonemes on the supercoiled DNA in the presence of ParB. **C)** Kymographs showing supercoiled DNA (top) and ParB^alexa647^ proteins (bottom) in the single-molecule assay. **D)** Quantification of supercoiled plectoneme DNA amount (top), its diffusion coefficient (middle), and ParB^alexa647^ intensity signal (bottom) over the time of the kymograph show in C). **E-H)** Same as A-D for 25 nM ParB concentration. **I-J)** Schematic representation of supercoiled condensate in the presence of multiple ParB proteins.

At high ParB concentrations, however, the behavior was strikingly different. All DNA supercoiling plectonemes were found to converge and pin at one spot, namely the position of the ParB-DNA condensate (Fig. 3E, F, Fig. S6A-F).No dynamic movement of the condensed spot along the DNA was observed (Fig. 3G), which contrasts the data on naked non-supercoiled DNA molecules where the condensate size remained highly variable (as shown previously in Ref. 15). Quantitative analysis of the amount of DNA within the main plectonemic cluster, showed an increase of the DNA amount within the cluster over time by 40% (from ∼8 kb in the plectonemes before condensation to ∼11.5 kb for the final condensate, Fig. 3H). The absence of any plectonemes outside the condensate indicates that the ParB-DNA condensate absorbed all supercoiling writhe into a cluster that we term a ‘supercoiled condensate’. This supercoiled condensate was a static object with a negligibly low diffusion coefficient (∼2 kb^2^/s) (Fig. 3H - middle). Overall, the presence of large numbers of ParB proteins on the DNA drastically changed the dynamics of the supercoiled DNA.

To quantify the effects of supercoiling on the ParB-DNA condensation, we screened a range of ParB concentrations (0.5 nM – 25 nM) on both negatively and positively supercoiled DNA (Fig. 4). We observed that the presence of DNA supercoils did not hinder ParB proteins from spreading all over the DNA molecule at any concentration (Fig. S7), in line with a recent *in vivo* study^58^. Both on non-coiled, negatively, and positively supercoiled DNA molecules, the ParB signal spanned over the entire length of the molecule due to diffusion (Fig. S7).

**Figure 4.**
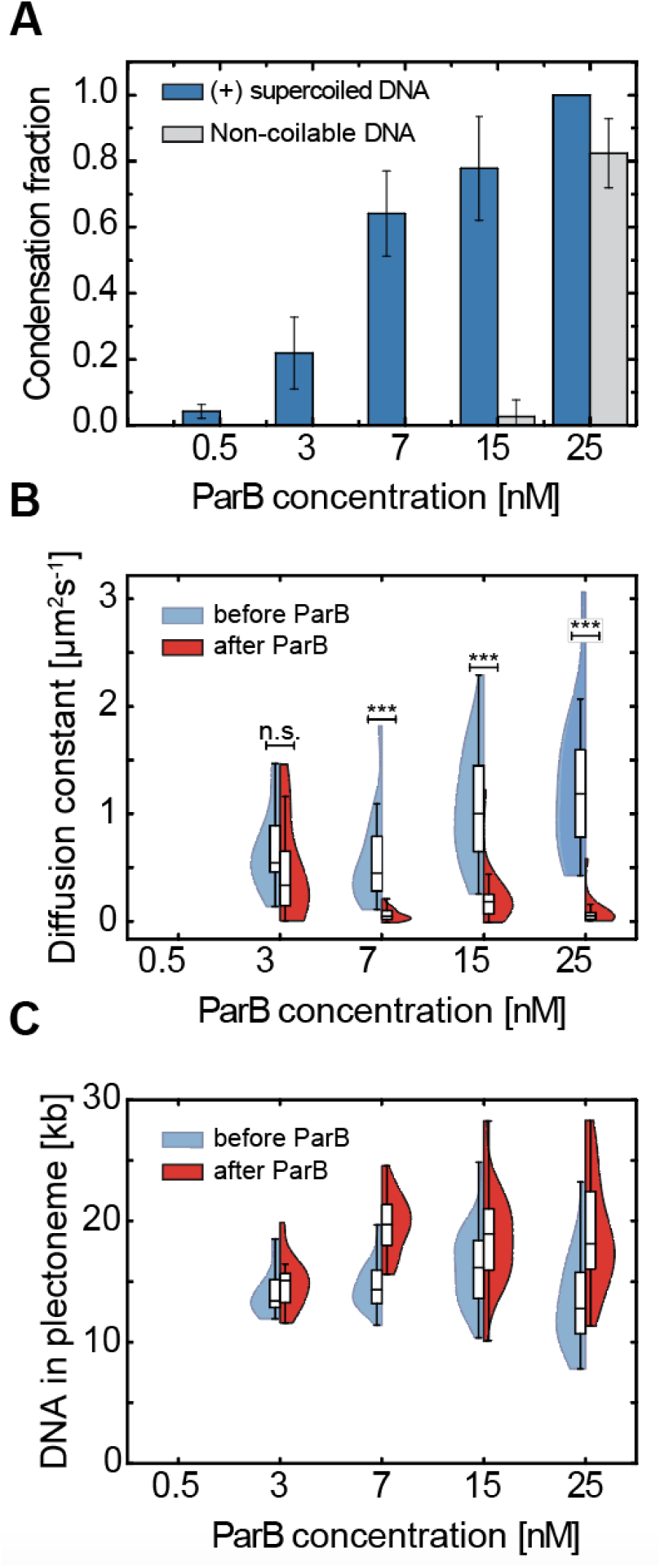
DNA-condensation by ParB proteins is facilitated by DNA supercoiling. **A)** Fraction of condensed molecules in the presence of an increasing concentration of ParB proteins on non-coiled (gray) and positively supercoiled DNA (blue). Error bars represent the binomial 95% confidence interval. (N - 0.5nM:0 (no condensed) 3nM:12 7nM:34 15nM:21 25 nM:16). **B)** Dynamics of DNA plectonemes/condensed plectonemes in the presence of increasing concentration of ParB proteins. Blue – before ParB enters the flow channel. Red – after >10 minutes after ParB was added to the flow channel and is covering the DNA*_parS_* molecules (p-values - 0.5nM: N/A; 3nM: 0.54; 7nM: <0.01; 15nM: 0.19; 25nM: 0.011). **C)** Total amount of DNA in plectonemes or in a supercoiled condensate on the 38 kb DNA*parS* molecules before (blue) and after (red) the addition of ParB at shown concentration.

An important observation was that the presence of DNA supercoiling decreased the minimal ParB concentration that is required for DNA condensation from ∼20 nM to ∼3 nM. This was observed for both positive (Fig. 4A) and negative supercoiling (Fig. S8A). While torsionally unconstrained DNA molecules did not form condensates at ParB concentrations below ∼25 nM, supercoiled DNA molecules (of either supercoiling sign) showed a sizeable fraction of molecules (∼25%) that exhibited condensation already at 3 nM. The one-dimensional diffusion constant of local DNA spots (i.e. plectonemes or supercoiled condensates at low and high ParB concentrations, respectively) showed a gradual reduction with increased concentrations of ParB (Fig. 4B). At the highest concentrations used (25 nM), almost all plectonemes pinned near the middle of the DNA molecules (around the *parS* site) with a diffusion constant close to zero (Fig. 4B). This strong reduction in the dynamics of plectonemes was found to be independent of the handedness of the supercoiling, as our data for negative supercoiling (Fig. S8B) showed the same phenomena as for positive supercoiling (Fig. 4B). When the condensed plectonemes pinned onto the DNA, they gradually increased the amount of DNA content within them over time (Fig. 3G, H). At the higher ParB concentrations in our experiments, we observed a sizable increase (∼40%) in the average DNA amount within the supercoiled condensate (Fig. 3C, S8C), albeit this varied between experiments.

### ParB condensate formation collapses linear extended plectonemes

While our data clearly show a pronounced DNA condensation of supercoiled DNA by ParB, the above experiments did not resolve much of the internal structure of the condensate, i.e., whether it is a globular condensed cluster or a linear extended object such as plectoneme. Current models for DNA condensation by ParB proteins propose stochastic bridging interactions of distant segments^12,15^ and possible DNA hairpins formed by laddering DNA segments^35^.

To resolve more of the internal structure, we used both our DNA stretching assay and atomic force microscopy (AFM). In our single-molecule visualization assay, we tethered the molecule, induced supercoiling, and induced ParB condensation, as in Fig. 1. In addition, however, we now exerted an in plane lateral flow that moved the molecule sidewards on the surface, revealing the inner topology of the DNA (Fig. 5A). In such experiments with torsionally unconstrained DNA, the lateral flow extended the molecule into a U-shaped arc in the direction of the buffer flow (Fig. 5B - left). When doing such experiments after addition of the ParB at high concentration (25 nM), however, the DNA molecules showed a high-intensity condensed spot near the middle of the DNA (Fig. 5B - right). This resembles the condensed structure of the partition complex, as reported by multiple works previously ^12,13,15,45,57^.

**Figure 5.**
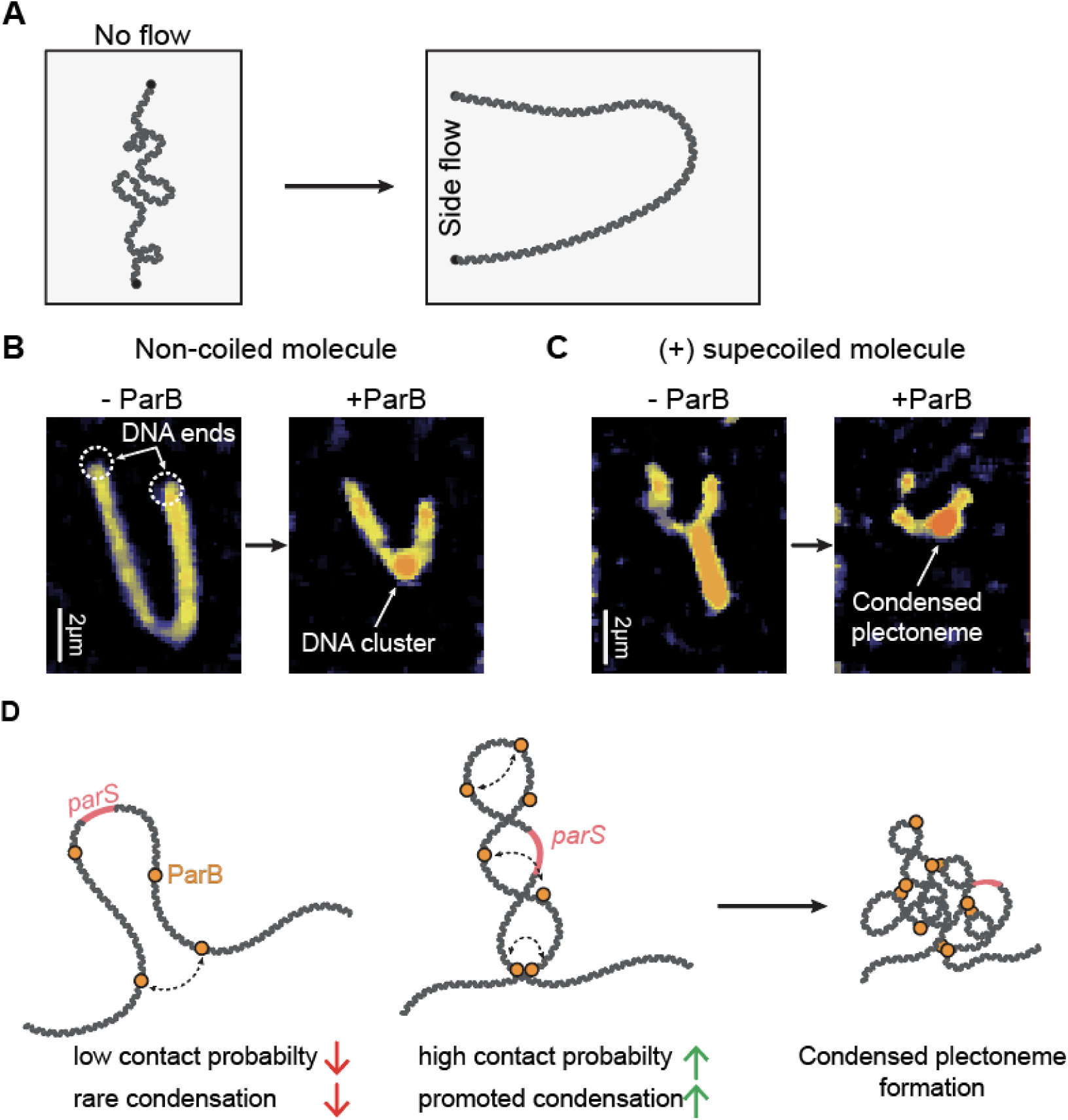
DNA plectoneme is condensed into a compact cluster in the presence of ParB. **A)** Schematic representation of the side-flow experiment. DNA is tethered parallel to the flow, then a lateral in-plane flow is applied that pushes the DNA in the U-shape only attached with its biotinylated DNA ends to the surface. **B)** Nicked DNA under side flow in the absence (left) and the presence (right) of 25 nM ParB. **C)** Same as B but for positively supercoiled DNA molecules. **D)** Sketches of the molecular conformations upon ParB-DNA condensation in the presence and absence of DNA supercoiling.

In the experiment with supercoiled DNA without ParB, the lateral flow caused instead one long plectonemic structure to align in the flow direction^38^ (Fig. 5C - left), which presumably results from the merging of multiple dynamic plectonemes into a single long plectoneme. After ParB was loaded on the supercoiled DNA under the same conditions (25 nM), however, the extended structure was found to have collapsed into a non-extended high-intensity spot near the middle of the DNA (Fig. 5C - right), which markedly differed from the previous extended plectoneme structure. In fact, the cluster resembled the same shape as the ones on the non-coiled DNA molecules. These images showed that for ParB on supercoiled DNA, the plectonemes were completely condensed into a compact ParB-DNA cluster that had absorbed all supercoiling writhe.

To observe the DNA structure below the optical resolution, we used AFM on 4.2 kb circular DNA*_parS_*that was either supercoiled or nicked. The nicked molecules showed typical open conformations on AFM surface (Fig. S9A). After the addition of ParB to these non-supercoiled DNA, we observed high compaction in large regions of the DNA molecule or even encompassing the entire DNA molecule (Fig. S9B) – in line with previous studies^12,15^. For supercoiled DNA molecules (without ParB), we observed extended DNA molecules with multiple crossings, typical of plectonemic DNA (Fig. S9C). Upon addition of ParB to supercoiled DNA molecules, the plasmids showed entirely condensed structures, where any plectonemic regions could not be resolved due to the high compaction and protein coverage in the ParB-DNA clusters (Fig. S9D). The AFM data confirm the observations of collapsed plectonemic structure in fluorescence assay experiments.

## Discussion

DNA supercoiling is an important regulator of many essential processes such as transcription, replication, and DNA compaction and segregation^26,59,60^. Vice versa, these processes, e.g. transcription by RNA polymerase induce supercoiling into the genomic DNA ^28,29,31^. As a result, bacterial DNA is continuously supercoiled^61,62^, both in the bacterial genome and in plasmid DNA. Both DNA supercoiling and the ParABS system promote distant intramolecular interactions by, respectively, plectoneme formation^33^ and condensate formation by ParB-ParB bridging^12,13,15,23,25,57^. In this work, we addressed the question of how these two processes affect one another.

### The impact of DNA supercoiling on ParB

We observed that the presence of DNA supercoiling did not alter ParB binding or condensate formation on the DNA molecules (Fig. 1-3). This finding is in line with recent *in vivo* data on plasmid DNA molecules^58^. Interestingly, ParB molecules bound efficiently onto negatively supercoiled DNA, non-coiled DNA, and positively supercoiled DNA, and all these cases could diffuse along these DNA substrates. No local pinning of plectonemes to locally bound ParB was observed. This can be attributed to the atypical topological binding of ParB to DNA: while most DNA-binding proteins firmly bind to a tight DNA-protein interface, ParB proteins only briefly interact with their *parS* recognition sequence ^55,63^, whereupon they release from it, while topologically encircling the DNA^6,11^. This topological entrapment of the DNA allows them to freely diffuse along DNA, irrespective of the sequence or twist of the DNA. We observed a somewhat faster diffusion but shorter residence times of ParB in the presence of DNA supercoiling, independent of its handedness (positive and negative). This increased one-dimensional diffusion may be due to an altered affinity of clamped ParB (i.e. after release from *parS*) to supercoiled DNA whereby it can slide faster on the DNA that contains twist. The decreased residence time is not trivial from the structural point of view, as the CTP binding pocket (N-terminus) of ParB is distant from the lumen that entraps the twisted DNA (C-terminus). While CTP-hydrolysis rates are unaffected, the faster release time of ParB may be due to a lowered affinity of the C-terminus to the supercoiled DNA backbone post-hydrolysis.

While DNA supercoiling does not greatly affect the behavior of single ParB proteins on DNA, we observed that it has a very strong effect on the DNA condensate that is formed by multiple ParB proteins. The concentration required to form a condensate on DNA was strongly (>5-fold) reduced in the presence of DNA supercoiling, which contrasts a previous report using magnetic tweezers^57^. Supercoiling thus greatly facilitates the partition complex formation. This may be attributed to increased intramolecular interactions between distant DNA segments, which occurs in plectonemic DNA, allowing ParB-ParB bridges to be formed with a higher probability (Fig. 5D). Similar observations were made in previous modelling studies that indicated that the introduction of DNA supercoiling promoted the formation of partition complexes ^35,36^. A recent study by Sekkouri Alaoui *et al.*^58^ however suggested no significant influence of supercoiling on the formation of partition complex in plasmids. This was concluded from ChIP-seq and fluorescence *in vivo* data, whereby it is difficult to deduce the three-dimensional structure of the ParB-DNA complex. Our data corroborate the findings that ParB can still efficiently spread over the supercoiled DNA (similar to ChIP-seq data^58^), independent of the supercoiling handedness, but we additionally show that the formation of the three-dimensional partition complex is strongly promoted by DNA supercoiling.

### The impact of ParB on supercoiled DNA

While supercoiling strongly affected the condensation of ParB proteins on DNA, ParB also drastically changed the dynamics of supercoiled DNA. As DNA condensation by ParB proteins proceeded, the motion of the rapidly moving plectonemes slowed down, until finally a single static spot on the DNA emerged (Fig. 3, 4). We termed this structure a ‘supercoiled condensate’ as it is a ParB-DNA condensate that absorbed all writhe, i.e., all plectonemic supercoils. This condensate had lost the characteristic linearly extended plectonemic structure (Fig. 5). A striking effect on the dynamics of DNA plectonemes was thus observed.

Interestingly, previous work showed that brief rifampicin treatment of bacterial cells showed a complete loss of any higher-order organization within the origin region^64–66^, which is surprising as ParB proteins should have been unaffected in their ability to locally condense the DNA in the partition complex. This observation is consistent with the hypothesis that supercoiling is a crucial facilitator for the maintenance of the ParB-DNA partition complex, especially considering the small number of ParB proteins (∼250- 700) in a bacterial cell^23,25^. In fact, supercoiling appears to underlie most of the large-scale chromosome compaction in bacterial cells^67^, and – as we show here – partition complex formation as well.

The ParABS system also facilitates the segregation and propagation of many plasmids in bacterial cells ^68,69^. Plasmids are often supercoiled due to continuous high expression of genes that enable their survival, and consecutive replication cycles ^70^, and for their small sizes (∼1-100kb^71^) supercoiling often has a significant effect on the entire molecule rather than on a local fraction. The effect of facilitated partition complex formation in the presence of DNA supercoiling therefore also has significant implications for plasmid biology.

Overall, this study provides key mechanistic insights into how two essential processes within the bacterial cells, DNA supercoiling and the DNA segregation machinery, interact.

## Methods

### ParB purification and fluorescent labeling

We prepared *Bacillus subtilis* ParB^L5C^ expression constructs using pET-28a derived plasmids through Golden-gate cloning. We expressed recombinant proteins in *E. coli* BL21-Gold (DE3) for 24 h in ZYM- 5052 autoinduction medium at 24°C. Purification of ParB^L5C^ variant, used for fluorescent labeling, was performed as described before^6,39^. Briefly, we pelleted the cells by centrifugation and subjected them to lysis by sonication in Buffer A (1 mM EDTA pH 8, 500 mM NaCl, 50 mM Tris-HCl (pH 7.5), 5 mM β-mercaptoethanol, 5 % (v/v) glycerol, and protease inhibitor cocktail (PIC, SigmaAldrich). We then added ammonium sulfate to the supernatant to 40% (w/v) saturation while stirring at 4°C for 30 min. We centrifuged the sample, collected the supernatant, and subsequently added ammonium sulfate to 50% (w/v) saturation and kept stirring at 4°C for 30 min. We collected the pellet (containing ParB^L5C^ proteins) and dissolved it in Buffer B (50 mM Tris-HCl (pH 7.5), 1 mM EDTA pH 8 and 2 mM β- mercaptoethanol). Before loading onto a Heparin column (GE Healthcare), the sample was diluted in buffer B to achieve a conductivity of 18 mS/cm. We used a linear gradient of buffer B containing 1 M NaCl to elute the protein. After collecting the peak fractions, we repeated the dilution in buffer B to 18 mS/cm conductivity and loaded it onto HiTrap SP columns (GE Healthcare). For elution, we used a linear gradient of buffer B containing 1 M NaCl. We loaded the collected peak fractions directly onto a Superdex 200-16/600 pg column (GE Healthcare) preequilibrated in 300 mM NaCl, 50 mM Tris-HCl (pH 7.5), and 1 mM TCEP. For fluorescent labeling, we incubated purified ParB^L5C^ variant with Alexa647-maleimide at a 1:2 molar ratio (protein:dye). We incubated the mixture for 15 min on ice, centrifuged it for 10 min, and then eluted it from a spin desalting column (Zeba) and flash frozen in liquid nitrogen. We estimated the fluorophore labelling efficiency at 74% for ParBalexa647 (resulting in 93% labelled) by an inbuilt function on Nanodrop using extinction coefficients of ε = 270 000 cm^-1^M^-1^ for Alexa647.

### Construction and purification of coilable 38 kb DNA*_parS_* construct for fluorescence experiments

To prepare a linear fragment adapted for flow cell experiments, we isolated ∼38 kb plasmid pBS-*parS* via a midiprep using a Qiafilater plasmid midi kit (Qiagen). The large plasmid was constructed from multiple smaller components as described in detail in Tišma *et al.* ^15^. We digested the pBS-*parS* for 2 h at 37°C using NotI-HF or XhoI restriction enzymes (New England Biolabs) and heat-inactivated for 20 min at 80°C. This resulted in the linear fragment that contains the *parS* site close to the middle of the DNA molecule, more specifically at the 0.4 relative position to the DNA ends. To prepare a linear fragment that would allow the introduction of DNA supercoiling upon the change of intercalating dye concentration, we constructed a fragment carrying handles with multiple biotins at the ends that would torsionally constrain the molecule from rotation around its axis. The handles for the 38 kb construct were made by PCR using primers CD21/CD22 of a 514 bp from the larger template pJT186 (see Table 2, and Tišma *et al.*^15^ for details and sequences) in the presence of 1:5 ratio of biotin-16-dUTP (Jena Bioscience, NU-803-BIO16-L) to dTTP (Thermo Fisher Scientific, 10520651). This allows stochastic, multiple insertions of biotinylated nucleotides into the final DNA*_parS_* ends. We digested these biotinylated PCR fragments using NotI-HF or XhoI for 2 h at 37°C which resulted in the handles of ∼250 bp in length. We mixed the digested handles with the large 38 kb fragment in 10:1 molar ratio (biotin-handles to DNA*_parS_*) and added T4 DNA ligase (New England Biolabs, M0202L) and 1 mM ATP for ligation. The ligation was set overnight at 16 °C and subsequently heat-inactivated the next day for 20 min at 65 °C. To remove the excess biotin handles from the large fragment we used ÄKTA pure, with a homemade gel filtration column containing approximately ∼46 ml of Sephacryl S-1000 SF gel filtration media (Cytiva), run with TE buffer with 150 mM NaCl_2_. The sample was run at 0.3 ml/min. The collected fractions stored at 4 °C, until use, in order to avoid freeze-thaw cycles that would introduce nicks into the DNA molecules. The final mixture contains ∼30-40% coilable molecules and the rest non-coilable/nicked, which served as the control comparison throughout this work.

**Table 2.**
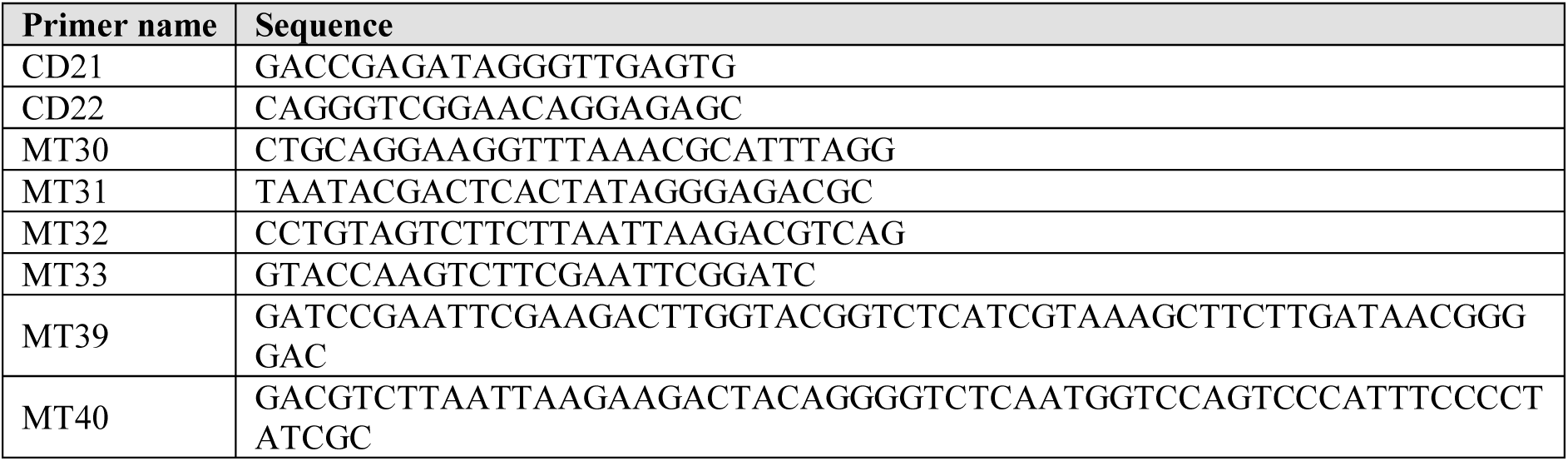
DNA primers used in the presented study for cloning and plasmid construction.

### Single-molecule visualization assay

We performed the experiments with supercoiled DNA and ParB proteins in custom-made flow cells, built by connecting a surface-passivated glass slide and a glass coverslip using double-sided tape^40,72^. The surfaces were prepared as described in detail by Chandranoss *et al.*^72^ with slight modifications. After extensive cleaning, the surface was silanized using APTES (10% v/v) and acetic acid (5% v/v) methanol solution. We passivated the surface with NHS-ester PEG (5,000 Da) and biotinylated NHS-ester PEG (5,000 Da) in relation ∼40:1. This step was repeated 4x24h to ensure low sticking of ParB proteins to the surface. Additionally, we treated the surface with 0,5 mg/ml UltraPure™ BSA (ThermoFisher Scientific), for 30 min in T20 buffer (40 mM Tris-HCl pH 7.5, 20 mM NaCl). This further reduced the non-specific sticking of labelled ParB to the surface.

For immobilization of 38 kb DNA*_parS_*, we introduced 50 μl of ∼3 pM of biotinylated-DNA*_parS_* molecules at a flow rate of 1.5 – 4 μl/min in imaging buffer without the oxygen scavenging enzymes (40 mM Tris-HCl, 2 mM Trolox, 1 mM TCEP, 30 mM Glucose, 2.5 mM MgCl_2_, 65 mM KCl, 0.25 mg/ml BSA, 1 mM CTP, 25-400 nM SxO)). Immediately after the flow, we further flowed 100 μl of the wash buffer (40 mM Tris–HCl, pH 7.5, 20 mM NaCl, 65mM KCl, 25-400 nM SxO)) at the same flow rate to ensure stretching and tethering of the other end of the DNA to the surface. By adjusting the flow, we obtained a stretch of around 20−50% of the contour length of DNA. The positive and negative supercoiling was induced by changing the SxO concentration during the initial tethering of the DNA*_parS_* and during imaging. Namely, to induce positive supercoiling of the tethered DNA, we tethered the DNA at the initial 25 nM SxO while the final experiments are done in 250 nM SxO in the imaging buffer (40 mM Tris-HCl, 65 mM KCl, 2.5 mM MgCl_2_, 1 mM CTP, 2 mM Trolox, 1 mM TCEP, 10 nM Catalase, 18.75 nM Glucose Oxidase, 30 mM Glucose, 0.25 µg/ml BSA, 50-250 nM SYTOX Orange (SxO, ThermoFisher Scientific)). Conversely, to induce negative supercoiling the initial tethering was done in 400 nM SxO while the final imaging is done at 50 nM SxO. The release of prebound SxO dyes after immobilization of the DNA results and now presence of the lower amount of dyes in the DNA results in negative supercoiling of the DNA.

Next, we flowed in the imaging buffer without ParB protein at a very low flow rate (0.2 µl/min) to enable minimal disturbances to the DNA molecules before and after protein addition. Real-time observation of ParB diffusion was carried out by introducing ParB (0.2 - 25 nM) in the imaging buffer. We used a home-built objective-TIRF microscope for fluorescence imaging. We used alternating excitation of 561 nm (0.2 mW) and 647 nm (14 mW) lasers in Highly Inclined and Laminated Optical sheet (HiLo) microscopy mode, to image SxO-stained DNA and alexa647-labelled ParB respectively. All images were acquired with an PrimeBSI sCMOS camera at an exposure time of 100 ms (10 Hz frame rate), with a 60x oil immersion, 1.49 NA CFI APO TIRF (Nikon).

### Image processing and analysis

Areas with single DNA molecules were cropped from the raw image sequences and analyzed separately with a custom-written interactive python software ^73,74^. For the analysis, we smoothened the cropped image section using a median filter with a window size of 10 pixels, and the subtracted the background with the “white_tophat” operation provided in the *scipy* python module^75^. We adjusted the contrast of obtained images manually for visualization purposes only (i.e., Fig. 3C). The ends of a DNA were manually marked. To get kymographs of our image sequences, we obtained total fluorescence intensity of 11 pixels across the axis of the DNA and stacked them over time axis(i.e., Fig. 2B). We chose the same DNA axis to obtain kymograph of the ParB fluorescence channel.

To further analyze the kymographs, we identified the peaks of high DNA and protein intensity using the *scipy* python module and merged into tracks. From these tracks, we calculated the size of a plectoneme or DNA condensate from the fraction of fluorescence intensity in the tracked peak relative to the overall fluorescence intensity of the DNA. We used an 11-frame moving window to calculate the apparent diffusion constant *D* over time (Fig. 2D, H). We calculated the diffusion constants of plectonemes (i.e., Fig. 4B) for each analyzed DNA molecule from the MSD calculated over a lag time ranging from 2 to 20 frames, then by fitting to the function *MSD* = 2*D*τ where *τ* is the lag time. The condensation fraction as reported in Fig. 4A was calculated from manual identification based on the presence of ParB-Alexa647, a high DNA signal, and a low apparent diffusion constant. As DNA plectonemes can be confused for condensed segments of DNA, we screened for both the presence of ParB and significant change in the plectoneme diffusion dynamics (D < 10 kb^2^/s) over an extended time period (200 frames, i.e. 40 sec) before classifying an event as ‘supercoiled condensate’ vs just a local diffusing plectoneme. The 1D curves diffusion signal, condensate size signal and ParB intensity signal (Fig. 3) were filtered using median filter of the window size of 9 frames. Analysis of plectoneme size vs DNA position as well as plectoneme position vs ParB cluster position was carried out using custom-written scripts in Igor Pro V6.39 (Wavemetrics, USA).

### Construction of 4.2 kb DNA*_parS_* construct for atomic force microscopy experiments

To construct the circular DNA for AFM experiments we used a commercially available pGGA plasmid backbone (New England Biolabs). We linearized the pGGA plasmid using MT032 and MT033 primers (Table 2). In parallel to this we extracted a region containing *parS* site downstream of *metS* gene in *B. subtilis* genome by a colony PCR using primers MT039 and MT040 (Table 2). We combined the plasmid backbone with the colony PCR insert by mixing them in molar ratio 1:3 in the 2xHiFi mix (New England Biolabs) to obtain the final plasmid of 4175 bp. We incubated the reaction at 50°C for 60 min, and cooled it down to 4°C for 30 min. We then transformed 2 µl of this reaction into 50 µl of *E. coli* NEB5alpha cells (New England Biolabs), and verified the presence of insert in grown colonies the following day by sequencing using MT030 and MT031 (Table 2). We grew sequence-positive clones for the plasmid extraction at 37°C overnight in the presence of a selective antibiotic Chloramphenicol (30 µg/ml). For obtaining supercoiled plasmids, we diluted the overnight culture 1:100 in 10 ml of fresh LB-Cm medium, and grew at 30°C until the culture reached OD_600_ = 0.6. We then placed the culture on ice for 5 min, and then spun down 4 ml of the culture before proceeding to isolation of the final plasmid using a QIAprep Spin Miniprep kit (Qiagen). The samples were stored at 4 °C, in order to avoid any freeze-thaw cycles that could introduce nicks into the DNA molecules and lower the yield of supercoiled plasmids. The plasmids that were going to be nicked were extracted directly from the overnight culture. These plasmids were nicked using a modified protocol for Nb. BbvCI nicking enzyme (New England Biolabs) capitalizing on a pre-existing recognition site in the *metS* gene. We mixed the solution containing the plasmids and the nicking enzyme was incubated at 37°C for 90 min and immediately purified over the PCR extraction membrane (Wizard® SV, Promega) and specifically skipped the recommended 80°C inactivation step, which would result in a fraction of single-stranded DNA in our AFM experiments. The following step was three rounds of plasmid clean-up using the same QIAprep Spin Miniprep kit (Qiagen) to remove all the residual enzymes that could corrupt the AFM images.

### Atomic force microscopy experiments and imaging

We obtained images in dry conditions using an AFM from Bruker Multimode 2 (Massachusetts, USA) and Scanassyst-Air-HR tips from Bruker. We incubated samples with different molarity ratios of DNA, CTP and ParB in Eppendorf tubes for 2 to 5 min in a buffered solution (40 mM Tris, pH 7.5, 70 mM KCl, 7.5 mM MgCl_2_). Then, we deposited the solution onto a freshly cleaved mica for 30 s. Afterward, we thoroughly washed the surface with 3 ml of Milli-Q water and dried it under a flow of Nitrogen until visibly dry. AFM was operated using peak force-tapping mode. We used WSxM software^76^ for all image processing and data extraction from our raw data in AFM experiments.

## Acknowledgments

We thank Jaco van der Torre for useful discussions on the analysis, data interpretation, and DNA constructs.

## Author contributions

Conceptualization: MT; Fluorescence experiments: AMG, MT; Formal analysis: AMG, BTK, RJ; Protein purification: HA; Atomic force microscopy experiments and analysis: AMG; DNA constructs and cloning: MT; Visualization: AMG, MT; Funding acquisition: SG, CD; Supervision: CD; Writing – original draft: MT; Writing – review & editing: all authors.

## Data and materials availability

All raw data from experiments are available upon request to corresponding authors.

