## Supplementary figures for "DNA supercoiling enhances DNA condensation by ParB proteins"

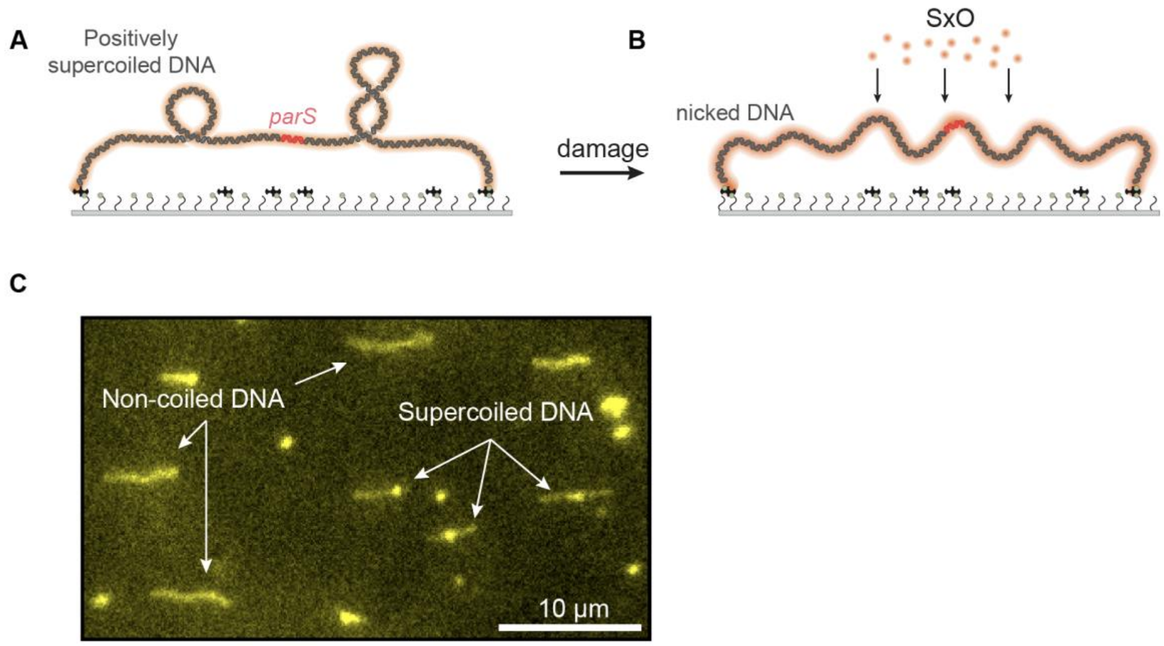


**Figure S1. *In vitro* single-molecule fluorescence detection of non-coiled DNA as well as plectonemes on supercoiled DNA. A)** Schematic representation of the single-molecule DNA stretching assay, with 38 kbp DNA$_{parS}$ tethered to the glass surface in the presence of SYTOX Orange intercalating dye. **B)** The same molecule if rendered non-coilable, which may e.g. occur due to nicking. Non-coiled DNA molecules appear brighter and more flexible. **C)** Typical section of the experimental field of view with both non-coilable and supercoiled molecules under the same conditions.

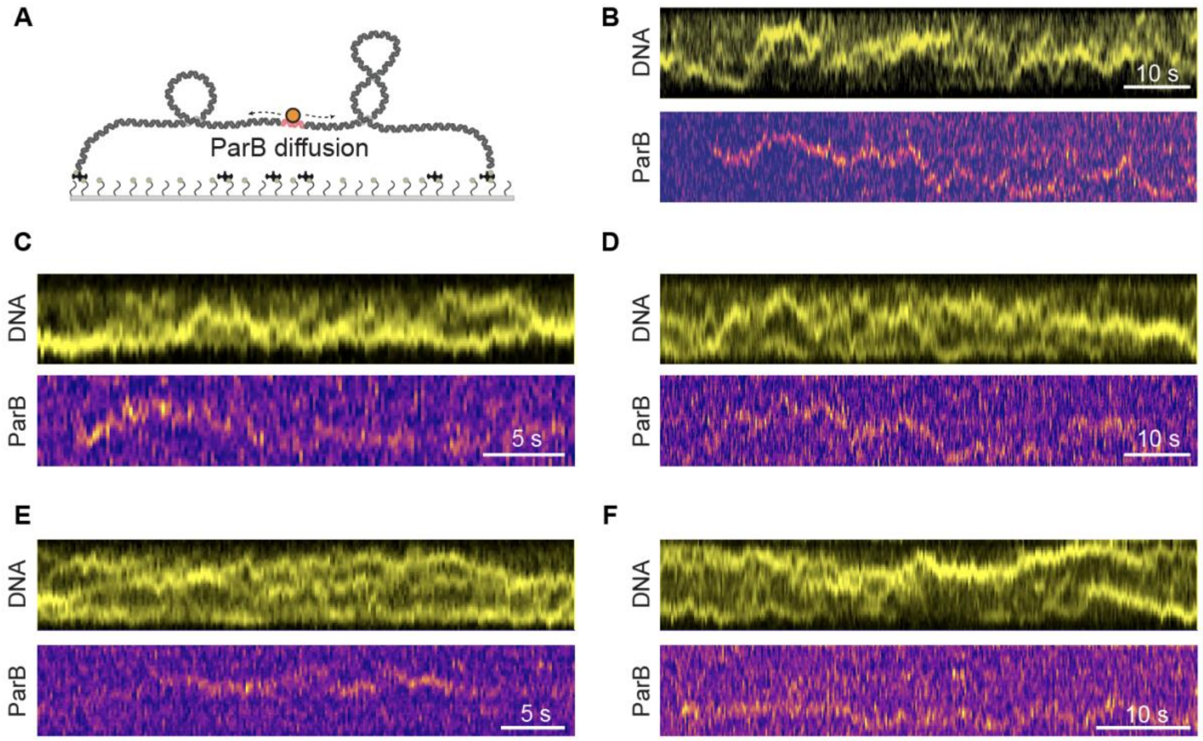


**Figure S2. ParB diffusing along a supercoiled DNA after binding the *parS* sequence.**
**A)** Schematic representation of the supercoiled DNA and diffusion of ParB proteins. **B-F)** Kymographs showing one-dimensional diffusion of a single $ParB^{alexa647}$ dimers (bottom) and DNA plectonemes (top) without strong plectoneme pinning.

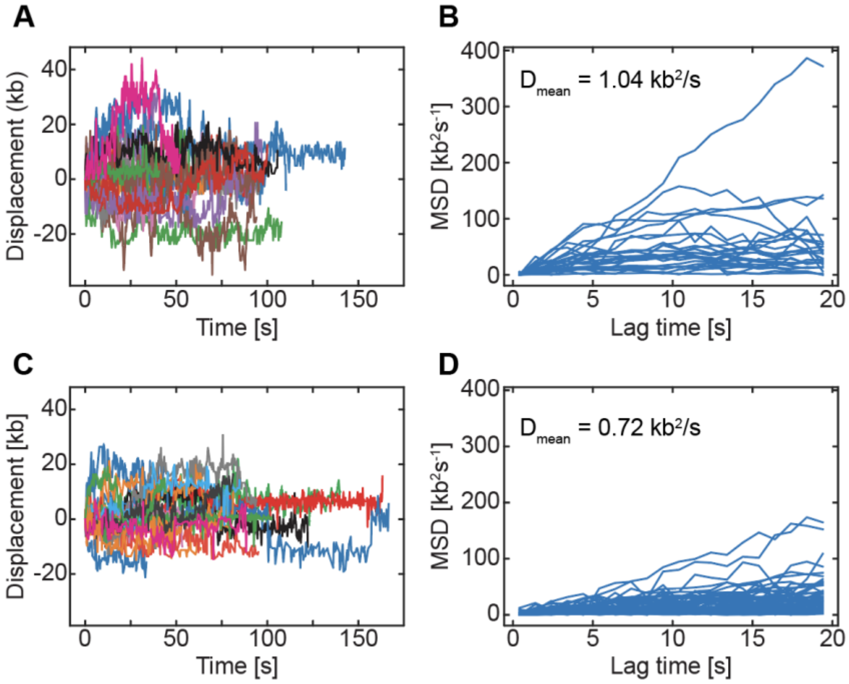


**Figure S3. ParB exhibits faster diffusion on positively supercoiled DNA.**
**A)** Single ParB diffusion traces along the 38 kb DNA molecule. **B)** Mean squared displacement (MSD) of individual ParB traces. $D_{mean} = 1.04 \pm 0.26$ $kb^2/s$ (mean ± SEM), $D_{median} = 0.69 \pm 0.35$ $kb^2/s$ (median ± SE), n = 66. **C-D)** Same as A-B but for non-coilable DNA molecules. **D)** $D_{mean} = 0.72 \pm 0.11$ $kb^2/s$ (mean ± SEM), $D_{median} = 0.43 \pm 0.13$ $kb^2/s$ (median ± SE), n = 58.

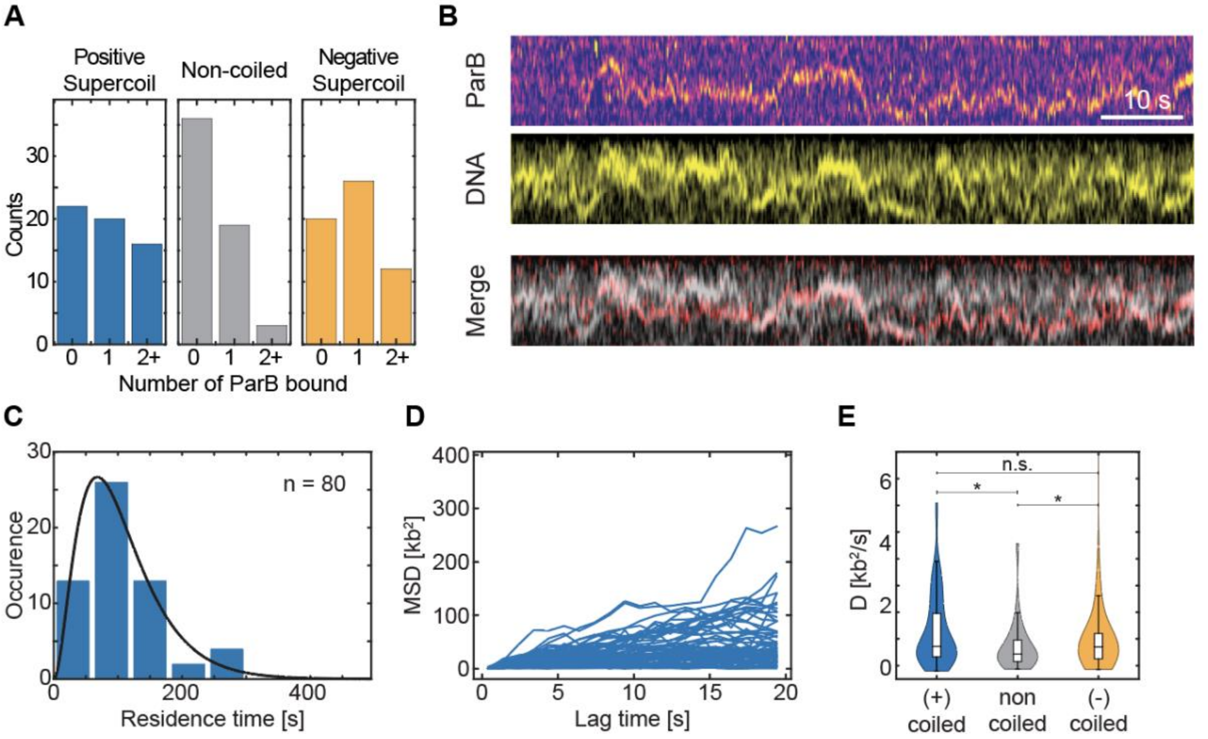


**Figure S4. ParB efficiently binds and diffuses along negatively supercoiled DNA. A)** Quantitative analysis of ParB molecules binding to positively coiled (Left), non-coiled (Middle) and negatively coiled (Right) DNA along 2000 frames. **B)** Kymographs showing one-dimensional diffusion of a single ParB$^{alexa647}$ dimers (top) and DNA plectonemes (middle). **C)** Residence times of diffusing ParB dimers after binding to the *parS* site. The data were fitted to a model assuming a delayed dissociation of ParB from the DNA after CTP hydrolysis of both nucleotides (black line, see Tišma et al. 2022). **D)** Mean squared displacement (MSD) of individual ParB traces on negatively supercoiled DNA. $D_{mean} = 1.31 \pm 0.20$ $kb^2/s$ (mean ± std), $D_{median} = 0.59 \pm 0.25$ $kb^2/s$ (median ± SE), n = 57. **E)** Diffusion coefficient of ParB molecules on non-coiled, negatively coiled and positively coiled DNA molecules (N: 58; 76; 70). P-values: p(negative-positive) = 0.14; p(negative-noncoiled) = 0.033; p(positive-noncoiled) = 0.014.

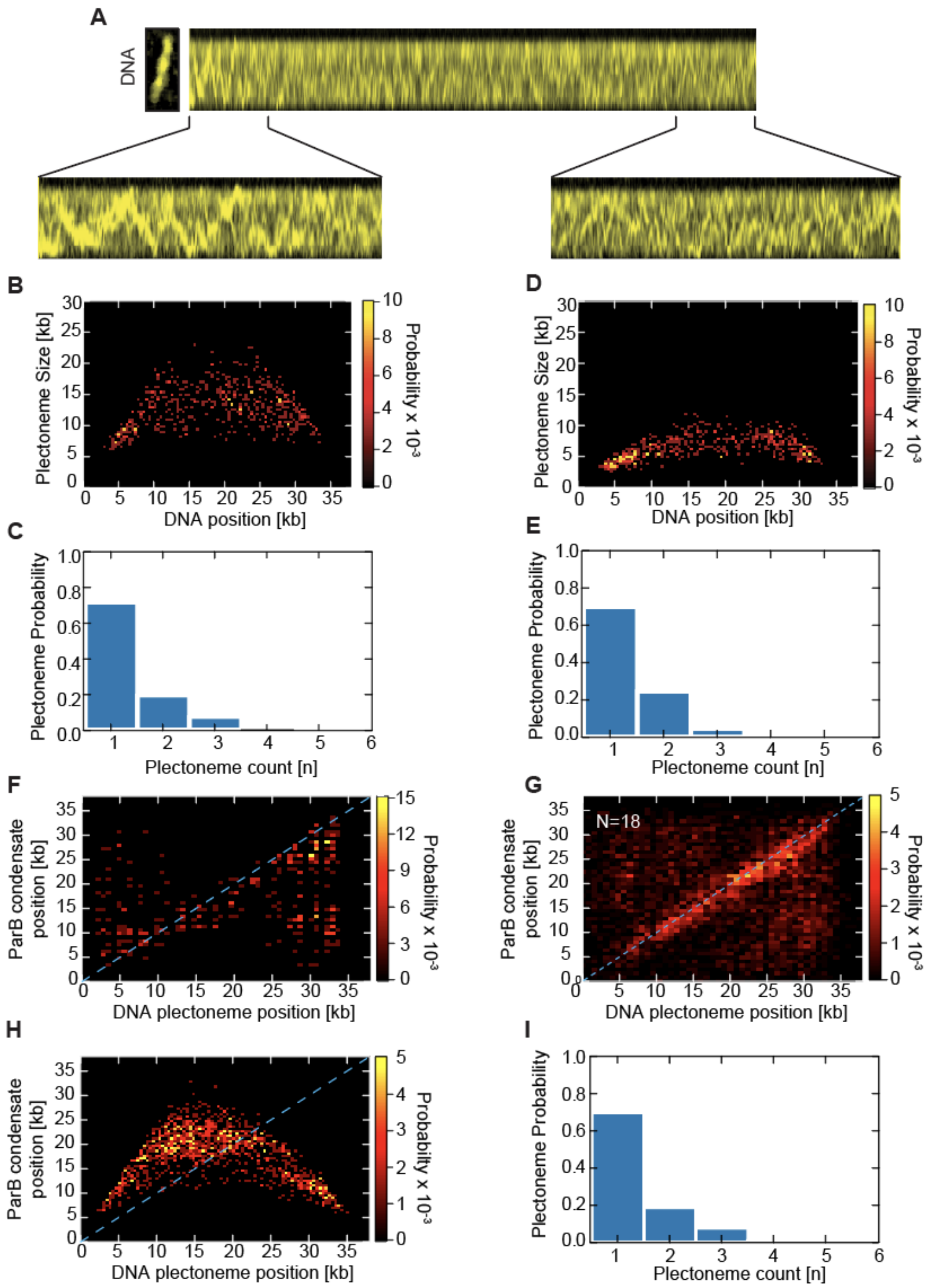


**Figure S5. ParB proteins at low concentration do not pin DNA plectonemes. A)** Kymographs showing supercoiled DNA at 3 nM [ParB] in the single-molecule assay. **B)** Probability distribution of plectoneme size versus DNA position before protein binding at 3 nM ParB protein. **C)** Observed number of co-existing plectonemes on the supercoiled DNA before the presence of ParB. **D)** Probability distribution of plectoneme size versus DNA position after protein binding at 3 nM ParB protein. **E)** Observed number of co-existing plectonemes on the supercoiled DNA in the presence of ParB. **F)** Heatmap showing the spatial correlation between ParB brightest condensate and brightest DNA plectoneme. **G)** Same as G) but cumulative for all molecules. **H)** Probability distribution of plectoneme size versus DNA position before protein binding at 0 nM ParB protein as a control. **I)** Observed number of co-existing plectonemes on the supercoiled DNA in the absence of ParB (show in G).

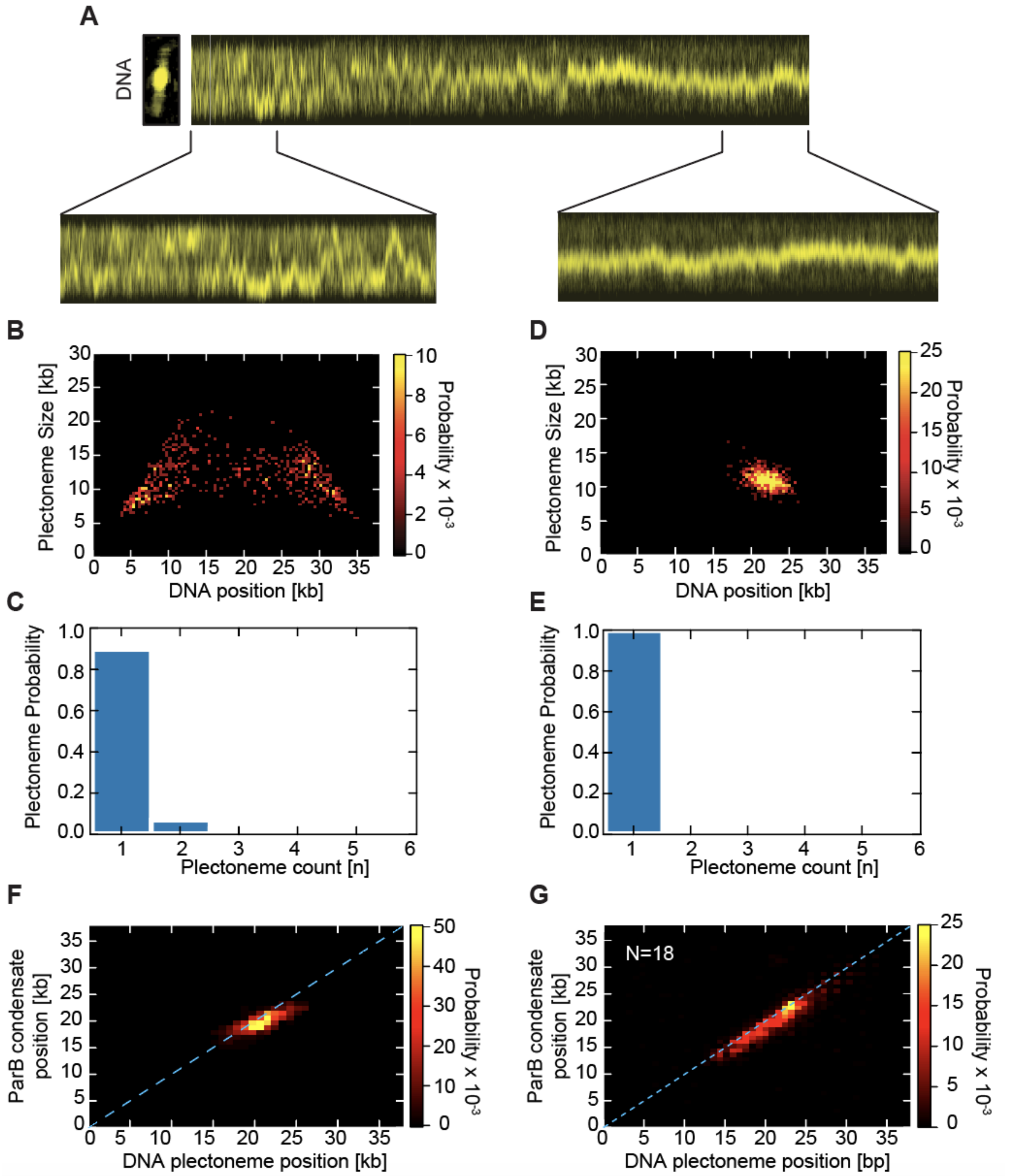


**Figure S6. Multiple ParB proteins pin the plectonemes into a single static cluster at high concentrations. A)** Kymographs showing supercoiled DNA at 25 nM [ParB] in the single-molecule assay. **B)** Probability distribution of plectoneme size versus DNA position before protein binding of ParB protein. **C)** Observed number of co-existing plectonemes on the supercoiled DNA before the presence of ParB (related to panel B). **D)** Probability distribution of plectoneme size versus DNA position after ParB protein binding at 25 nM concentration. **E)** Observed number of co-existing plectonemes on the supercoiled DNA in the presence of ParB (related to D). **F)** Spatial correlation between ParB brightest condensate and brightest DNA plectoneme. Blue line indicates the line of perfect correlation (r = 1). **G)** Same as in panel F but cumulative for all molecules (N=18).

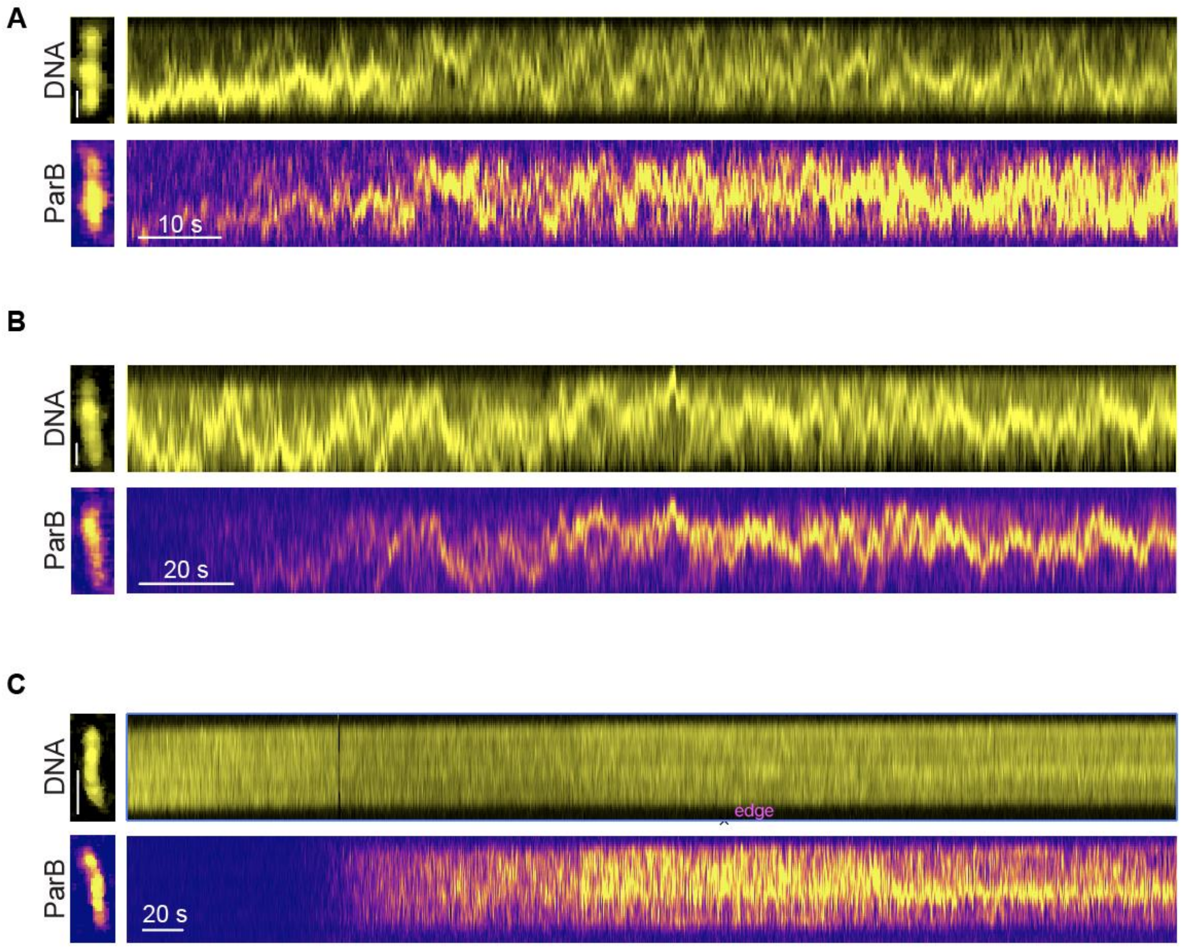


**Figure S7. ParB spreading is not affected by DNA supercoiling.**

**A-C)** Kymographs of ParB spreading over the entirety of the 38 kb DNA molecule at the concentration of 25nM. **A)** ParB spreading on positively supercoiled DNA. Spatial scale bar = 1 µm **B)** ParB spreading on negatively supercoiled DNA. Spatial scale bar = 1 µm. **C)** ParB spreading on non-coiled DNA molecule. Spatial scale bar = 2 µm.

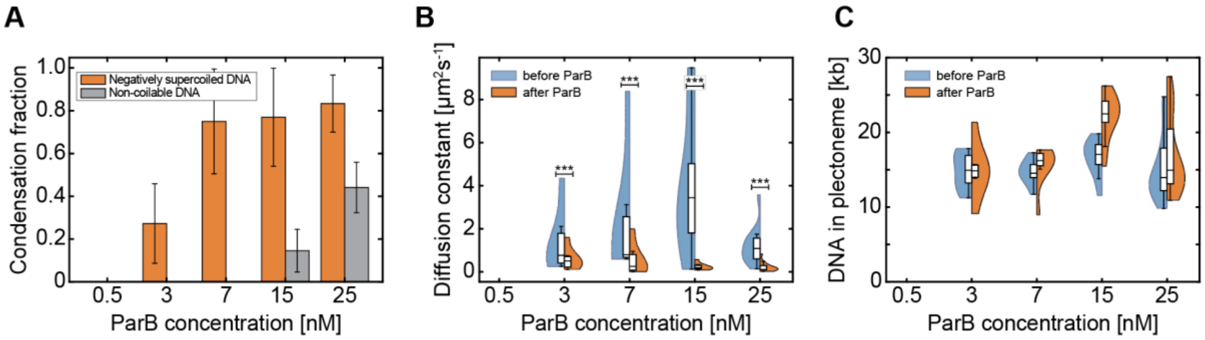


**Figure S8. DNA-condensation by ParB proteins is facilitated by DNA negative-supercoiling.**

**A)** Fraction of condensed DNA molecules in the presence of an increasing concentration of ParB proteins on non-coiled (orange) and negatively supercoiled DNA (blue). (N - 0.5nM:16; 3nM:22; 7nM:18; 15nM:17; 25nM:30) **B)** Dynamics of DNA plectonemes/condensed plectonemes in the presence of increasing concentration of ParB proteins. Blue – before ParB enters the flow channel. Orange – after >10 minutes after ParB was added to the flow channel and is covering the $DNA_{parS}$ molecules (p-values: 0.5nM:N/A; 3nM:0.93; 7nM:0.31; 15nM:<0.001; 25nM:<0.001). **C)** Total amount of DNA in plectonemes or in a negatively supercoiled condensate on the 38 kb $DNA_{parS}$ molecules before (blue) and after (orange) the addition of ParB at shown concentration.

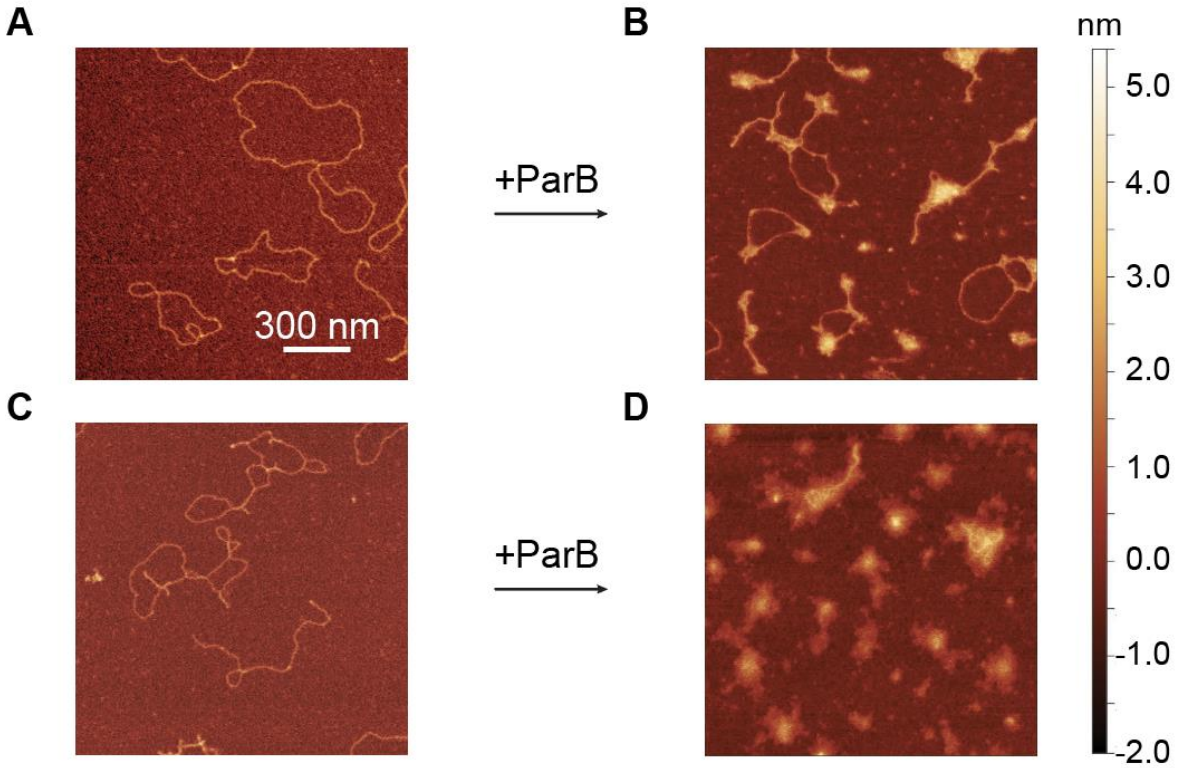


**Figure S9. ParB proteins collapse the extended structures of DNA plectonemes.**

**A)** AFM image of a nicked DNA plasmid with a length of 4.2 kb. **B)** AFM image of a nicked DNA incubated with 15 nM of ParB for 5 minutes leading to the formation of small DNA-condensates. **C)** Same as A but the plasmid was maintained in a supercoiled state. **D)** AFM image of a supercoiled DNA molecule incubated with 15 nM of ParB for 5 minutes leading to the collapse of extended DNA plectonemes into condensed structures.
